# PUPpy: a primer design pipeline for substrain-level microbial detection and absolute quantification

**DOI:** 10.1101/2023.12.18.572184

**Authors:** Hans Ghezzi, Michelle Y. Fan, Katharine M. Ng, Juan C. Burckhardt, Deanna M. Pepin, Xuan Lin, Ryan M. Ziels, Carolina Tropini

## Abstract

Characterizing microbial communities at high-resolution and with absolute quantification is crucial to unravel the complexity and diversity of microbial ecosystems. This can be achieved with PCR assays, which enable highly selective detection and absolute quantification of microbial DNA. However, a major challenge that has hindered PCR applications in microbiome research is the design of highly specific primer sets that exclusively amplify intended targets. Here, we introduce Phylogenetically Unique Primers in python (PUPpy), a fully automated pipeline to design microbe- and group-specific primers within a given microbial community. PUPpy can be executed from a user-friendly GUI, or two simple terminal commands, and it only requires coding sequence files of the community members as input. PUPpy-designed primers enable the detection of individual microbes and quantification of absolute microbial abundance in defined communities below the strain level. We experimentally evaluated the performance of PUPpy-designed primers using two bacterial communities as benchmarks. Each community was comprised of 10 members, exhibiting a range of genetic similarities that spanned from different phyla to substrains. PUPpy-designed primers also enable the detection of groups of bacteria in an undefined community, such as the detection of a gut bacterial family in a complex stool microbiota sample. Taxon-specific primers designed with PUPpy showed 100% specificity to their intended targets, without unintended amplification, in each community tested. Lastly, we show absolute quantification of microbial abundance using PUPpy-designed primers in ddPCR, benchmarked against 16S rRNA and shotgun sequencing. Our data shows that PUPpy-designed microbe-specific primers can be used to quantify substrain-level absolute counts, providing more resolved and accurate quantification in defined communities than short-read 16S rRNA and shotgun sequencing.

**Importance:** Profiling microbial communities at high resolution and with absolute quantification is essential to uncover hidden ecological interactions within microbial ecosystems. Nevertheless, achieving resolved and quantitative investigations has been elusive due to methodological limitations in distinguishing and quantifying highly related microbes. Here, we describe PUPpy, an automated computational pipeline to design taxon-specific primers within defined microbial communities. Taxon-specific primers can be used to selectively detect and quantify individual microbes and larger taxa within a microbial community. PUPpy achieves substrain-level specificity without the need for computationally intensive databases and prioritises user-friendliness by enabling both terminal and graphical user interface (GUI) applications. Altogether, PUPpy enables fast, inexpensive, and highly accurate perspectives into microbial ecosystems, supporting the characterization of bacterial communities in both *in vitro* and complex microbiota settings.

## Introduction

Microbial communities play a crucial role in numerous aspects of life on Earth, including nutrient cycling, disease development, and carbon fixation (1, 2). These processes are driven by a remarkable genetic and functional diversity spanning various taxonomic levels, from unrelated phyla to closely related species. Although functional diversity is greatest among unrelated microbes, it is also present in microorganisms with closely related genetic makeups. Importantly, subtle genetic variations at the strain level can also lead to extensive phenotypic differences (3, 4), such as changes in microbial metabolism (5–7), immunomodulation (8, 9), antibiotic resistance (10, 11), biofilm formation (12), and pathogenesis (13). For example, the impacts of different *Escherichia coli* strains on human health can vary significantly: Nissle 1917 (14) is commensal, while enterohemorrhagic *E. coli* (EHEC) O157:H7 and enteroaggregative *E. coli* (EAEC) O104:H4 (15) are pathogenic. Thus, profiling microbial communities with at least strain-level granularity is imperative to uncover hidden ecological interactions that may otherwise be missed at lower taxonomic resolution. However, detecting and quantifying closely related microbes remains a challenging task due to their largely indistinguishable genetic makeup, often limiting microbial profiling to less-resolved taxonomic levels.

Over the last decade, advances in both culture-dependent and independent approaches have enabled increasingly high-throughput and high-resolution characterization of microbial communities. In the case of culture-dependent methods, while significant progress in growing bacteria in controlled environments has enabled the identification of bacterial strains and species from mixed communities, the work-intensive nature of these methods has limited their widespread application (16, 17). On the other side, culture-independent methods, specifically amplicon (e.g., 16S rRNA) and shotgun sequencing have enabled cost-effective and high-throughput detection and quantification of microbes, including rare and unculturable ones (18). Advances in both short and long-read technologies, as well as profiling methods, have significantly improved taxonomic resolution, which remains lower in 16S rRNA than shotgun sequencing due to the reduced variability in the amplicon sequenced (19, 20). In addition, several approaches have been employed to estimate absolute abundance from sequencing data, such as spike-ins, cell counts, and DNA concentration (21, 22). While these strategies provide an estimate, accurately quantifying absolute abundance remains an inherently challenging task due to numerous factors, including difficulties in quantifying the microbial load contribution for each microbe present, contamination of plant or host DNA, loss of DNA during extraction, and distinct biases uniquely affecting different DNA fragments at multiple steps of the sequencing workflow (21–23).

Another method for granular microbial detection and absolute quantification involves selectively amplifying microbe-specific genes. This can be achieved through widely available polymerase chain reaction (PCR) techniques, such as quantitative PCR (qPCR) and droplet digital PCR (ddPCR) (24, 25). These methods are ubiquitously established, cost and time-effective, and highly sensitive, facilitating resolved and targeted microbial investigations without considerable computational and monetary resources. Furthermore, the sensitivity of PCR assays enables exceptionally accurate absolute quantification of microbial populations, including rare ones. Nevertheless, designing highly selective primer sets at any taxonomic resolution (e.g., genus, species, or strain level) remains a major challenge that has hindered the application of PCR assays in microbial investigations. This becomes especially arduous when surveying complex and undefined microbial communities (e.g., the fecal microbiome), where the genetic makeup of each microbe is not known *a priori*. Commonly, primer design was focused on universal phylogenetic markers such as the 16S rRNA amplicon and housekeeping genes, such as *rpoB* (*26*)*, gyrB* (*27*), and *tuf* (*28*). However, these molecular markers often lack the discriminatory power to detect closely related microbes due to the limited variations in the amplicon considered. More recently, advances in sequencing technologies have yielded an outstanding increase in assembled microbial genomes, thus enabling the design of unique phylogenetic markers across the entire genetic material and increasing the resolution achievable.

Currently, several tools exist to design primers specific to input genomic sequences, including SpeciesPrimer (29), find_differential_primers (fdp) (30), RUCS (31), and TOPSI (32). However, these tools have either been discontinued, require significant manual handling to generate configuration files, or cannot flexibly design both microbe- and group-specific primers for any user-defined combination of microbes. Another previously published primer design toolset is DECIPHER (33), which offers multiple versatile functionalities focusing on databases, including primer design with several *in silico* PCR steps. However, this tool requires coding expertise to manually parse the databases over numerous commands, which limits its usability. In addition, users must manually provide previously aligned DNA sequences as input, further adding complexity to primer design. Altogether, these challenges highlight the need for a user-friendly, streamlined, high-throughput, and highly selective taxon-specific primer design tool.

Here, we introduce Phylogenetically Unique Primers in python (PUPpy), a fully automated pipeline to design primers targeting individual microbes or groups of microbes within a given microbial community. PUPpy only requires coding sequences (CDSs) of the microbes of interest as input to design primers with substrain level specificity. The pipeline streamlines primer design by only requiring 2 commands, which can be run either from the terminal or from an intuitive graphical user interface (GUI). We benchmarked PUPpy-designed primers against two defined bacterial communities, each consisting of 10 members with varying degrees of genetic similarity, ranging from distinct phyla to substrains. We also evaluated the ability of PUPpy-designed primers to detect members of a highly diverse and fastidious gut bacterial family, the Muribaculaceae, in a complex and undefined conventional mouse microbiota. We show that all taxon-specific primers designed with PUPpy selectively amplified the respective targets in all communities tested. Finally, we assessed the effectiveness of PUPpy-designed primers for accurately quantifying the total microbial count in specific communities. This assessment was conducted using ddPCR and was compared with results obtained from 16S rRNA and shotgun sequencing methods. Our data suggests that taxon-specific primers enable more resolved and accurate quantification than short-read 16S rRNA and shotgun sequencing in defined microbial communities. Altogether, PUPpy represents an intuitive tool that enables absolute quantification of individual microbial taxa, an essential parameter that has been missing in microbiome studies. The versatility of PUPpy makes it suitable for a variety of applications involving both defined and complex communities, ranging from environmental studies to host-associated microbiota research.

## Results

### Overview of the PUPpy pipeline

PUPpy is a fully automated computational pipeline that allows the design of taxon-specific PCR primers targeting user-defined microbial communities (Fig. 1A). This pipeline defines two primary applications of taxon-specific primer design: microbe- and group-specific (Fig. 1B). Microbe-specific primers target unique CDSs only present in a single member of the community. These primers allow users to target individual taxa with full specificity, a crucial aspect in tracking and quantifying specific microbes in communities with high degrees of genetic similarity. Conversely, group-specific primers are designed to select genes that are shared across all user-defined targets (Fig. 1B). Importantly, these primers enable the selective assessment of microbial dynamics of an entire taxon (e.g., all *Bacteroides*) in a sample. In both microbe and group-specific applications, PUPpy designs taxon-specific primers by aligning every CDS in the community and identifying unique (microbe-specific) or shared (group-specific) genes. Because PUPpy only requires CDS files as input, it can run locally in most cases, and does not require memory intensive databases (e.g., the NCBI nr database). More details on the computational workflow are provided in the Methods section. To demonstrate the applicability of PUPpy in microbiome research, we empirically validated taxon-specific primers in 3 increasingly complex microbial communities. These communities were composed of 1) a defined community of 10 phylogenetically distinct bacteria, 2) a defined community of 10 bacteria from 3 increasingly related taxa, and 3) a complex microbiota from conventional mouse fecal samples. By gradually increasing the genetic similarity and complexity of the community, we were able to assess the resolution limits and effectiveness of the pipeline under diverse microbial community scenarios.

**Fig. 1.**
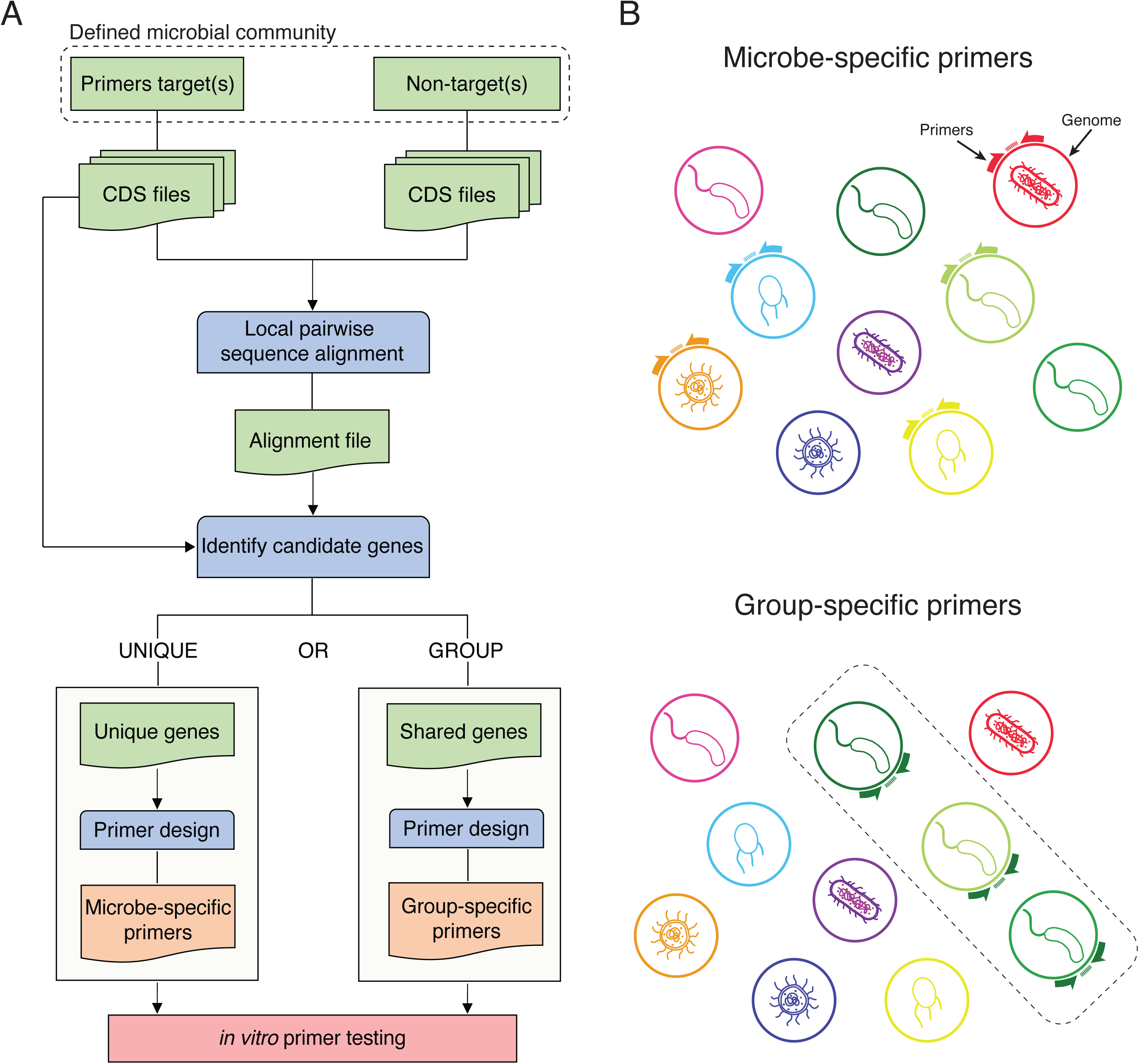
The PUPpy pipeline designs taxon-specific primers in defined microbial communities. (A) Overview of the PUPpy workflow. Input coding sequence (CDS) files for primer target(s) and non-targets (the background) are aligned using MMseqs2 (48). Candidate unique or shared genes are selected to design microbe- and group-specific primers, respectively, with Primer3 (62). (B) The key output of PUPpy is taxon-specific primers, which include microbe- and group-specific primers. Microbe-specific primers selectively target individual members of the community, while group-specific primers target user-determined collections of microbes.

### Microbe-specific primers show specificity to intended targets in a genetically diverse community

The first community, hereafter referred to as the “GUT community”, was composed of 10 phylogenetically distinct bacteria belonging to 9 families and 5 phyla (Bacillota, Bacteroidota, Pseudomonadota, Actinomycetota, Verrucomicrobiota) (Table S1). These bacteria were chosen as being phylogenetically distinct, common gut commensals, and representative of the 5 most abundant phyla of the human gut microbiota. This community, while simple, accurately represents a defined microbial consortium similar to that represented in several gnotobiotic animal models (34–36). Notably, each member of this community possesses a significant number of unique genes and has been previously characterized (34, 37, 38). To confirm the genetic diversity within the GUT community, we calculated Average Nucleotide Identity (ANI) on the genomic assemblies for the 10 bacteria (see Methods). The community included two *Bacteroides* species, *Bacteroides ovatus* and *Bacteroides thetaiotaomicron*, which, as expected, shared a greater degree of genetic similarity, with 80% percentage identity and ∼50% percentage coverage (Fig. 2A). All other GUT members displayed greater genetic diversity, with <75% percentage identity and <8.5% percentage coverage (Fig. 2A). Following ANI analysis, we ran PUPpy with default parameters and generated microbe-specific primers for all 10 GUT community members (see Table S2). Consistent with the degree of genetic diversity observed in the ANI analysis, PUPpy identified fewer unique CDSs in more genetically similar organisms. PUPpy orders primer targets based on descending number of unique genes found, enabling an immediate visual and quantitative evaluation of community diversity (Fig. 2B). Specifically, compared to all other community members, PUPpy identified ∼27% of the CDSs as unique in *B. ovatus* and ∼26% in *B. thetaiotaomicron,* the most related microbes, as opposed to ∼74% in *E. coli* BW25113 (Fig. 2B). Importantly, PUPpy considers all unique CDSs to design microbe-specific primers, and multiple primers can be designed on the same CDS. Thus, even in the organism with the fewest unique genes, *B. thetaiotaomicron*, PUPpy evaluated 1248 options (i.e., ∼26%) for primer design (Fig. 2B). Next, we empirically validated the specificity of the GUT microbe-specific primers by PCR and gel electrophoresis using gDNA isolated from the 10 GUT community members (see Methods). Each primer pair was tested against i) the respective gDNA positive control, ii) the 10-member gDNA pool, iii) a 9-member gDNA pool including all taxa except the primer target, and iv) a no-template control (Fig. 2C). Following this experimental validation, we found that all microbe-specific primers were specific to the respective primer target, without unintended amplification (Fig. 2D, S1). PUPpy can therefore be leveraged to design microbe-specific primers and selectively detect individual microbes in a genetically diverse and defined microbial community.

**Fig. 2.**
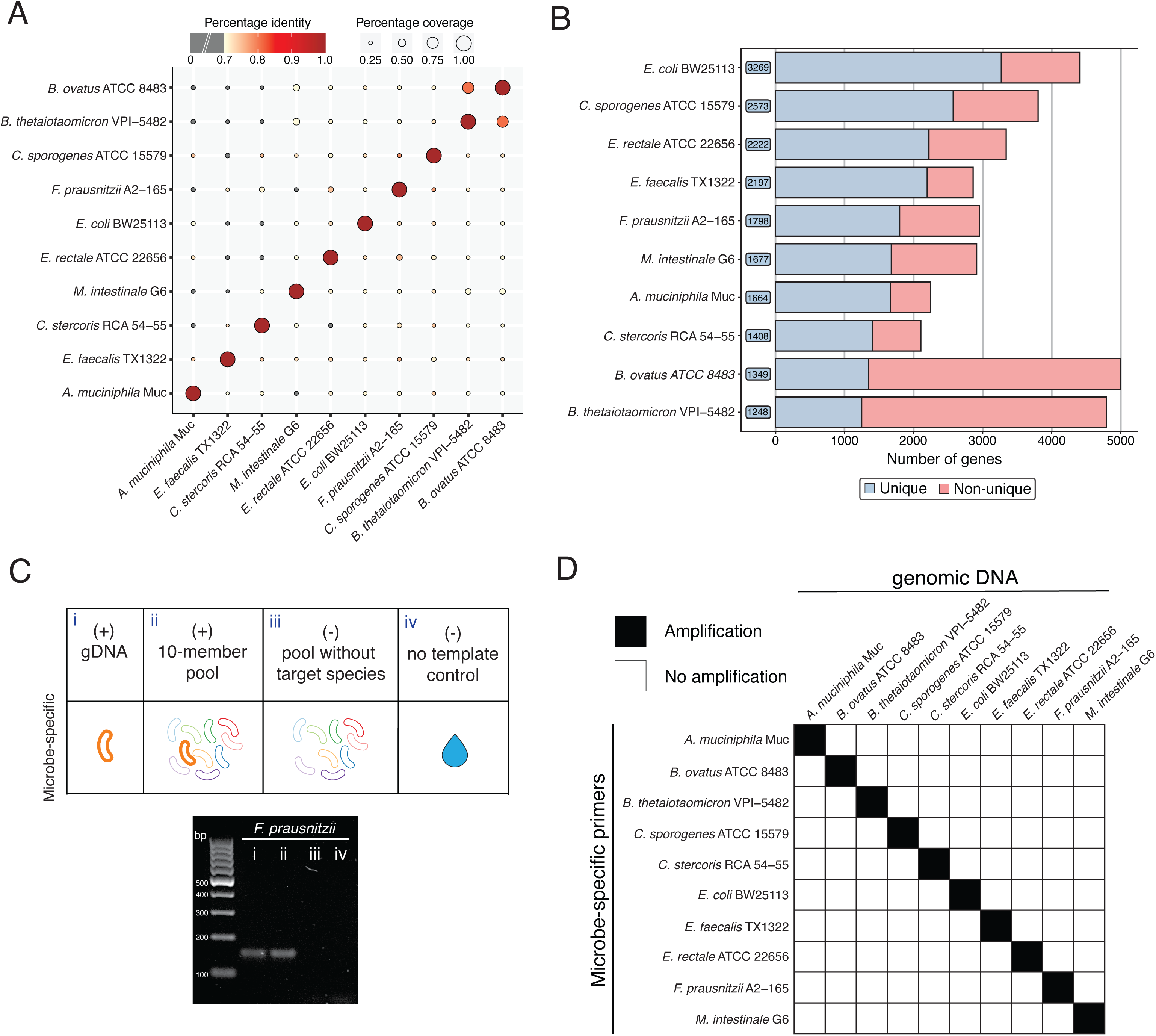
PUPpy-designed microbe-specific primers selectively detect microbes in a community of 10 phylogenetically distinct bacteria. (A) Pairwise Average Nucleotide Identity (ANI) based on the whole-genome sequences of the 10 GUT community members (Methods). Color indicates percentage identity, with darker red indicating greater similarity between sequences. Grey circles indicate percentage identity lower than 70%. Circle size indicates alignment coverage between sequences, or the percentage of 2 sequences (in this case genomes) that are aligned. (B) PUPpy-generated bar plot showing the number of unique genes (to the right of the strain name) identified for each member of the input community. The high number of unique genes found across GUT community members, ranging from 1248 in *B. thetaiotaomicron* (∼26% of the CDSs) to 3269 in *E. coli* BW25113 (∼74% of the CDSs), reflects the genetic diversity observed by ANI (A). (C) Experimental conditions for validation of microbe-specific primers specificity in the GUT community. The PCR gel image exemplifies the specificity of microbe-specific primers tested using the conditions above for *F. prausnitzii*. (D) Binary heatmap showing the targets amplified by each microbe-specific primer pair following the validation in (C). Each primer pair exclusively amplified the respective intended target (black squares along the diagonal) following primer optimization. Individual gel imaging data can be found in Fig. S1.

### Taxon-specific primers selectively detect microbes down to the substrain level

As microbial communities often harbor multiple species and strains within the same genus, we next asked whether PUPpy could be used to design taxon-specific primers and detect highly related microbes. To test this, we created a second 10-member bacterial community, hereafter referred to as the “Species-Strain-Substrain (SSS) community” (Table S1). This community consisted of 3 *Enterocloster* species, 3 *B. thetaiotaomicron* strains and 4 *E. coli* K-12 substrains. These bacteria were chosen to assess the taxonomic resolution achievable by PUPpy, while also maintaining a degree of phylogenetic diversity across taxa. To evaluate the genetic similarity in the community we performed ANI analysis, which grouped the SSS community members into 3 distinct memberships consistent with their assigned taxonomic classifications (Fig. 3A). As expected, the *Enterocloster* species displayed the lowest similarity, with a minimum of ∼78% identity and ∼34% coverage. Conversely, the closely related *E. coli* substrains were more genetically similar, sharing >99.95% identity and ∼99% coverage (Fig. 3A). Next, we ran PUPpy with default parameters to design both microbe-specific primers for each member and group-specific primers for each taxon (Table S2). Group-specific primers are designed on genes shared across all primer targets (e.g., all *Enterocloster* species) but missing in all unintended targets (e.g., all *B. thetaiotaomicron* strains and *E. coli* substrains), enabling the selective detection of microbial taxa within a microbial community. Consistent with the ANI analysis and taxonomic classification, PUPpy ordered the 10 members into 3 groups based on the number of unique CDSs found (Fig. 3B). Specifically, PUPpy identified between ∼19% and ∼36% unique CDSs for the 3 *Enterocloster* species, between ∼7% and ∼17% for the *B. thetaiotaomicron* strains, and between 0% and ∼1% for the *E. coli* K-12 substrains (Fig. 3B). Using default parameters, PUPpy did not identify any unique CDSs for *E. coli* MC4100 within the SSS community, and thus no microbe-specific primers were designed or tested for this organism (Fig. 3B). The lack of unique genes is likely due the close genetic similarity to all 3 other *E. coli* members. Increasing the percentage identity threshold in ‘puppy-align ‘may yield unique genes for *E. coli* MC4100 by decreasing the alignment stringency (see Methods for details). We validated the specificity of SSS microbe-specific primers following the same experimental design as the GUT community (Fig. 3C). In addition, we tested the specificity of group-specific primers by running each pair against i) the respective primer gDNA targets, (condition (ii) was not run as it would provide redundant information), iii) the 9-member SSS gDNA pool including all taxa except the target strain gDNAs, and iv) a no-template control (Fig. 3C). All microbe- and group-specific primers tested showed specificity to their intended targets, independent of taxonomic level and genetic similarity (Fig. 3C, 3D, S2). These data show that PUPpy enables the selective detection of microbes down to the substrain level by identifying and designing PCR primers for unique CDSs. In addition, this validation highlights the ability of PUPpy to identify conserved CDSs across specific groups of bacteria, allowing the selective amplification of multiple organisms with a single primer pair.

**Fig. 3.**
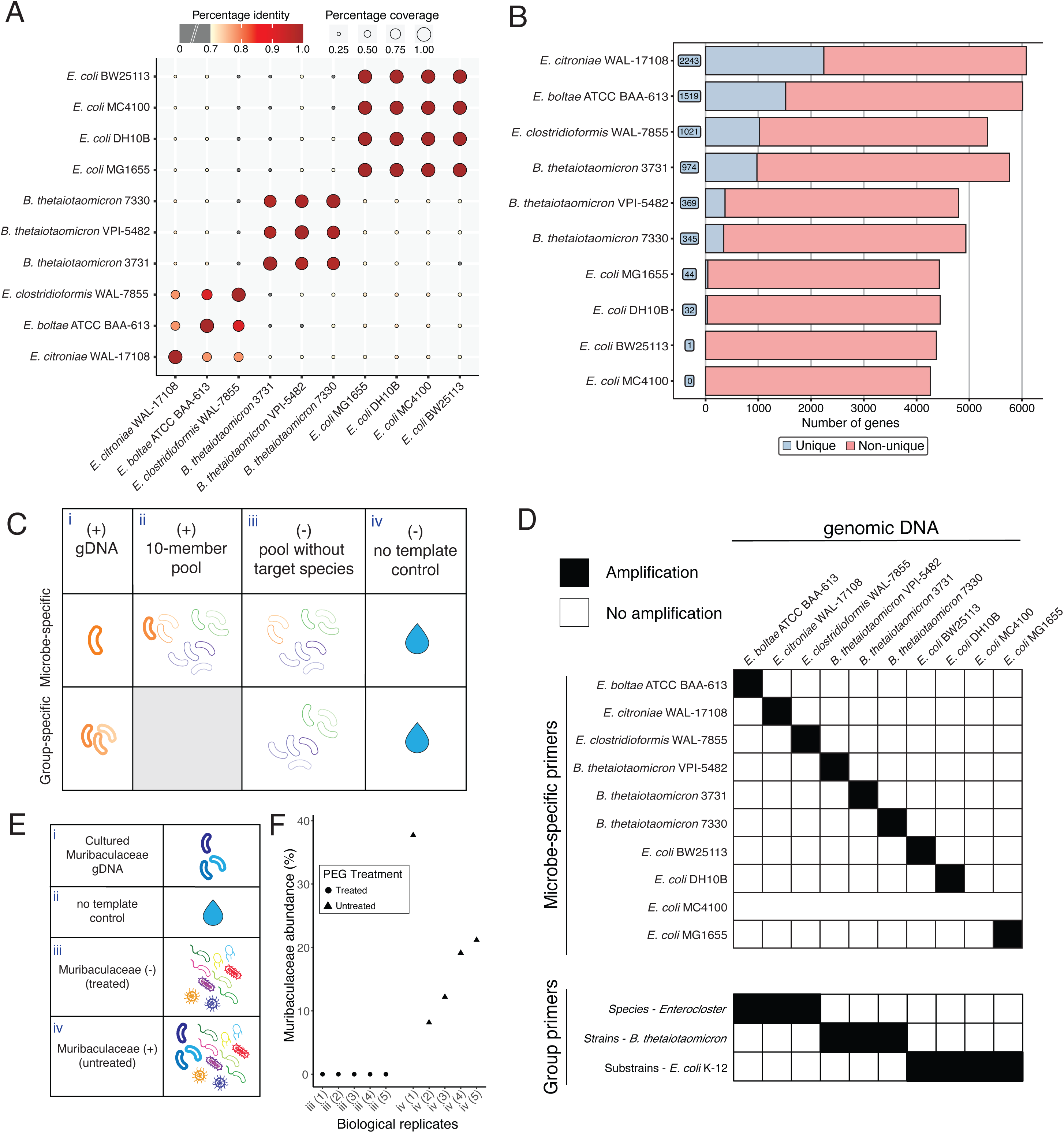
PUPpy-designed taxon-specific primers selectively detect microbes down to the substrain level. (A) Pairwise Average Nucleotide Identity (ANI) based on the whole-genome sequences of 10 microbes from 3 increasingly related taxa in the Species-Strain-Substrain (SSS) community. Color indicates percentage identity, with darker red indicating greater similarity between sequences. Grey circles indicate percentage identity lower than 70%. Circle size indicates alignment coverage between sequences, or the percentage of 2 sequences (in this case genomes) that are aligned. ANI analysis, assessing genetic similarity based on percentage identity and coverage, grouped the *Enterocloster* species, *Bacteroides thetaiotaomicron* strains, and *Escherichia coli* K-12 sub-strains as 3 distinct clusters consistent with their taxonomic assignment. *Enterocloster* species displayed lower genetic similarity, with a minimum of ∼78% identity and ∼34% coverage, while *E. coli* sub-strains were almost genetically identical, sharing >99.95% identity and ∼99% coverage. (B) PUPpy-generated bar plot. The number of unique genes identified by PUPpy in SSS community members decreases across increasingly related taxa. Between 19 and 36% of the CDSs in *Enterocloster* species are unique within the SSS community, while only 0-1% of CDSs are unique across *E. coli* K-12 sub-strains. (C) Experimental conditions for validation of microbe- and group-specific primers specificity in the SSS community. Group-specific primers were not validated against the 10-member community (i.e., experimental condition ii). (D) Binary heatmap showing selective amplification by both microbe- and group-specific primer pair down to the sub-strain level. Microbe-specific primer pairs could not be designed and validated for *E. coli* K-12 MC4100 because PUPpy could not identify unique genes for this organism. Individual gel imaging data can be found in Fig. S2. (E) Experimental conditions for validation of Muribaculaceae-specific primer specificity in a complex microbial community from conventional mouse fecal samples. (F) Muribaculaceae quantification by 16S rRNA sequencing of 5 Muribaculaceae-depleted (PEG-treated) (iii) and 5 Muribaculaceae-positive (untreated) (iv) biological replicates. Relative abundance was calculated using QIIME2 (69) (see Methods section).

### Group-specific primers selectively detect targets in a complex microbial community

Having validated the specificity of PUPpy-designed primers in defined microbial communities, we asked whether the pipeline could also be applied to complex communities in which the exact membership is not known. To investigate whether we could selectively identify the presence of specific taxa in this setting, we leveraged a conventional mouse gut microbiota. The mouse microbiota has high abundance of the family Muribaculaceae, a highly diverse and hard-to-culture bacterial family that is broadly prevalent in warm-blooded animals (39–41). We have previously shown that mice treated with the osmotic laxative polyethylene glycol (PEG) are depleted of Muribaculaceae (42) and we wanted to ascertain whether group-specific primers to this family could identify the loss of this family without the need for sequencing. We extracted gDNA from fecal samples of mice that were either untreated (Muribaculaceae-positive) or PEG-treated (Muribaculaceae-depleted) (see Methods). We confirmed Muribaculaceae presence or absence by performing 16S rRNA sequencing on the fecal gDNA samples, and then evaluated PUPpy-designed primers against the Muribaculaceae family. Such experimental validation presents two main challenges in the design of specific primers: 1) the overall composition of the community was treated as unknown *a priori* and therefore we did not have a complete list of CDS files to input into PUPpy and 2) we did not know which Muribaculaceae strains were present in the community of untreated mice, and thus for which microbes to design primers. Therefore, to minimize the chances of off-target amplification we selected a comprehensive list of 156 unintended targets and 11 intended Muribaculaceae targets (Table S1). The unintended targets were selected among prevalent gut commensal bacteria to represent members of the phyla Bacillota, Bacteroidota, Pseudomonadota, Actinomycetota, Verrucomicrobiota, and Fusobacteriota. Using this 167-member input community we were able to identify a gene shared across all Muribaculaceae but absent in all other 156 members. Importantly, since Muribaculaceae is a highly diverse family (40), we designed 8 primer pairs to account for higher potential representation of different sub-taxa in the complex community and pooled them to create a Muribaculaceae-specific primer cocktail. We validated this mix against both Muribaculaceae-positive and Muribaculaceae-depleted fecal samples and evaluated its specificity (Fig. 3E) (see Methods). Our results show that Muribaculaceae-specific primers selectively detected Muribaculaceae in the positive samples but did not in the depleted samples (Fig. S3), consistent with the results from 16S sequencing (Fig. 3F). In addition to being specific, these Muribaculaceae-specific primers did not display any off-target amplification of PCR amplicons with different sizes in the untreated samples (see Fig. S3). This is crucial in PCR assays such as qPCR and ddPCR, where quantification may be biased by unintended amplification. Altogether, we define a methodology involving the use of PUPpy to design taxon-specific primers that can selectively amplify microbial targets even in a complex community.

### Taxon-specific primers enable high resolution absolute microbial quantification

Beyond the detection of presence and absence of microbes, taxon-specific primers can also be used to quantify absolute microbial counts at extremely high-resolution using qPCR or ddPCR (24, 29). To evaluate microbial quantification with PUPpy, we compared quantification of PUPpy-designed primers via ddPCR to short-read 16S rRNA and shotgun sequencing. We pooled 10 extracted gDNA samples for each member of the GUT and SSS communities, respectively, in a theoretical 1:1 genomic copy ratio calculated from the respective microbial gDNA concentration and genome size (see Methods). We then aliquoted the gDNA pools for analysis with 16S rRNA sequencing, shotgun sequencing, and ddPCR with PUPpy primers (Fig. 4A) (see Methods). Using ddPCR we calculated absolute microbial abundance, converted it to compositional data, and compared it to the relative abundance estimated by the sequencing methods. In the GUT community, ddPCR and shotgun sequencing achieved similar resolution, which, as expected, was markedly higher than 16S rRNA sequencing due to the limited variability in the V3-V4 regions targeted in this technique (Fig. 4B). Specifically, amplicon sequencing classified only 4 members at the species level (*Akkermansia muciniphila, Eubacterium rectale*, *B. ovatus* and *B. thetaiotaomicron*) and the remaining 6 members at the genus level (Fig. 4B). Conversely, both shotgun sequencing and ddPCR identified all 10 members at the greatest taxonomic resolution. In addition, ddPCR provided a more quantitative assessment of bacterial composition by measuring absolute microbial counts, which is inherently more accurate than relative abundance.

**Fig. 4.**
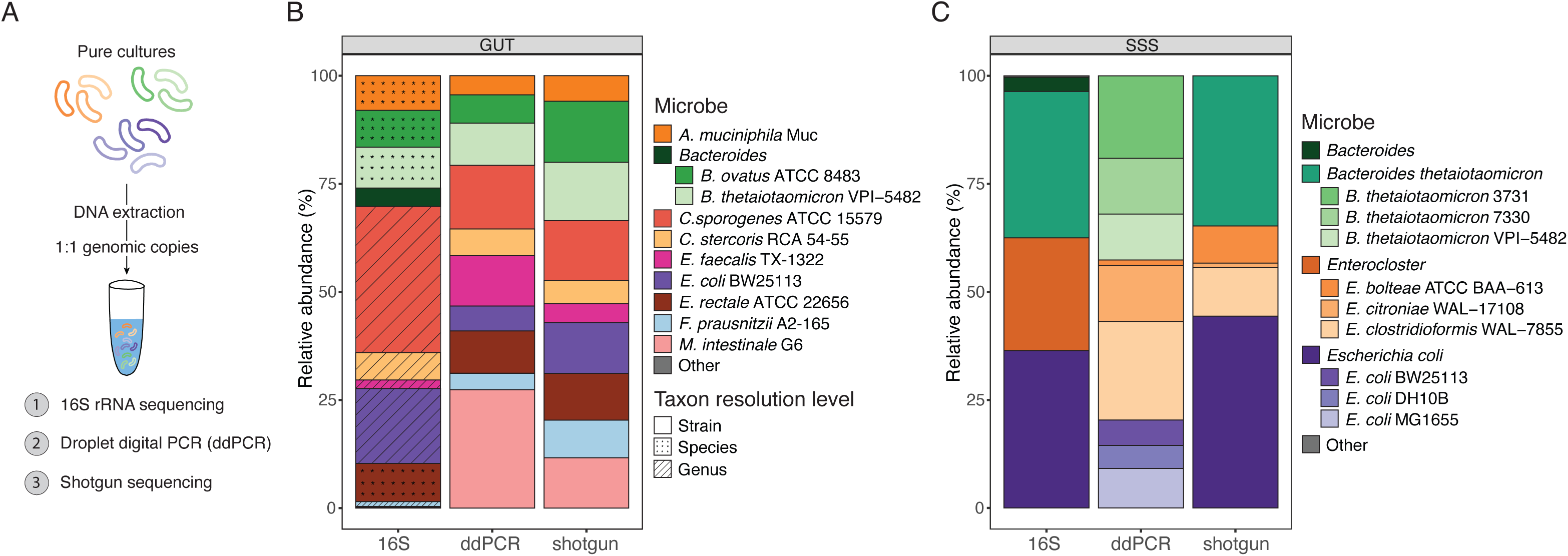
PUPpy-designed taxon-specific primers enable more accurate and resolved quantification than 16S and shotgun sequencing in defined communities. (A) Schematic of the workflow to prepare samples for quantification (see Methods section for details). (B, C) Stacked bar plots evaluating microbial quantification with droplet digital PCR (ddPCR), 16S and shotgun sequencing in the GUT (B) and SSS (C) communities. ddPCR absolute counts were converted to relative abundance for a more direct comparison to the other methods (see Methods).

ddPCR also yielded accurate and absolute quantification in the genetically related SSS members (Fig. 4C). This assay selectively detected and quantified absolute counts for all SSS community members, except *E. coli* K-12 MC4100, for which we did not design primers given its similarity within the community. Similar to the results for the GUT community, 16S rRNA sequencing achieved the lowest resolution, only detecting the 3 major taxa and no individual strains, while shotgun sequencing detected all 10 microbes (Fig. 4C). Although shotgun sequencing identified all individual members, its quantification accuracy was impacted by the high degree of genetic similarity within the community (Fig S4). In these scenarios, samples contain a substantial proportion of ambiguous reads (reads that map equally well to multiple targets in the reference database). Such reads can be either tossed out, assigned to the Lowest Common Ancestor (LCA), or randomly distributed across multi-mapping microbes (43). The fate of ambiguous reads varies with the goal of the study, pipelines used, and downstream applications. In the SSS community, the LCA approach would not achieve the resolution desired, and tossing out reads removed ∼60% of them, heavily underestimating microbial abundance in strains and substrains (Fig S4). Thus, we instructed Bowtie2 (44) to randomly distribute ambiguous reads and quantified microbes with coverM (45), which yielded a near uniform quantification of strains and substrains (Fig. 4C). However, we found this to be an artefact of randomly assigning multimapping reads, which coincidentally matched the expected input and would not extend to scenarios where strains and substrains are present in uneven proportions.

To further investigate the impact of multi-mapping reads for communities of arbitrary microbial concentrations, we generated *in silico* reads for the SSS members at known input proportions and analysed these synthetic reads using both the heuristic alignment approach used in Fig. 4 and Kraken2 (46), an LCA approach (see Methods for details). Unlike the heuristic alignment approach, this approach does not attempt to randomly assign ambiguous reads to multimapping targets, and instead assigns them to the LCA. We did not apply Bracken (47) to Kraken2 data to avoid collapsing quantification to the species-level, which would reduce the resolution needed to quantify the *B. thetaiotaomicron* strains and *E. coli* substrains (Fig S4). Both the heuristic alignment and LCA methods accurately quantified the 3 *Enterocloster* species, confirming the accuracy of shotgun sequencing when microbes are sufficiently distinct for reads to be confidently assigned (Fig. 5A and 5B). However, quantification accuracy with both pipelines gradually decreased as genetic similarity increased. Notably, the observed abundance obtained with the LCA approach for all strains and substrains was considerably lower than with heuristic alignment, consistent with the random distribution of ambiguous reads by the latter method (Fig. 5A and 5B). To further support this, we found that quantification with Kraken2 was considerably more accurate at the LCA level as opposed to the microbe level for both strains and substrains (Fig. 5C, 5D, and S4). Specifically, the ratio of observed vs expected relative abundance, expected to be 1 in perfect scenarios, was ∼0.80 for the *E. coli* LCA (Fig. 5C) while it dropped to between ∼0.05 and ∼0.001 for individual *E. coli* substrains (Fig. 5D). These data suggest that most reads belonging to closely related strains could not be unambiguously mapped with the methods tested and were randomly distributed to multi-mapping targets instead. These findings thus confirm that the shotgun sequencing quantification pipeline used in Figure 4 was not actually able to accurately quantify individual strains and substrains in the SSS community and instead yielded a uniform abundance by randomly distributing ambiguous reads (Fig. 4C). Altogether, these results shine light on the possible pitfalls of short-read sequencing and highlight the need for careful considerations of the pipelines and parameters used. Additionally, these results show that PUPpy-designed taxon-specific primers enable accurate detection and absolute quantification in defined microbial communities, even in the presence of highly related microbes.

**Fig. 5.**
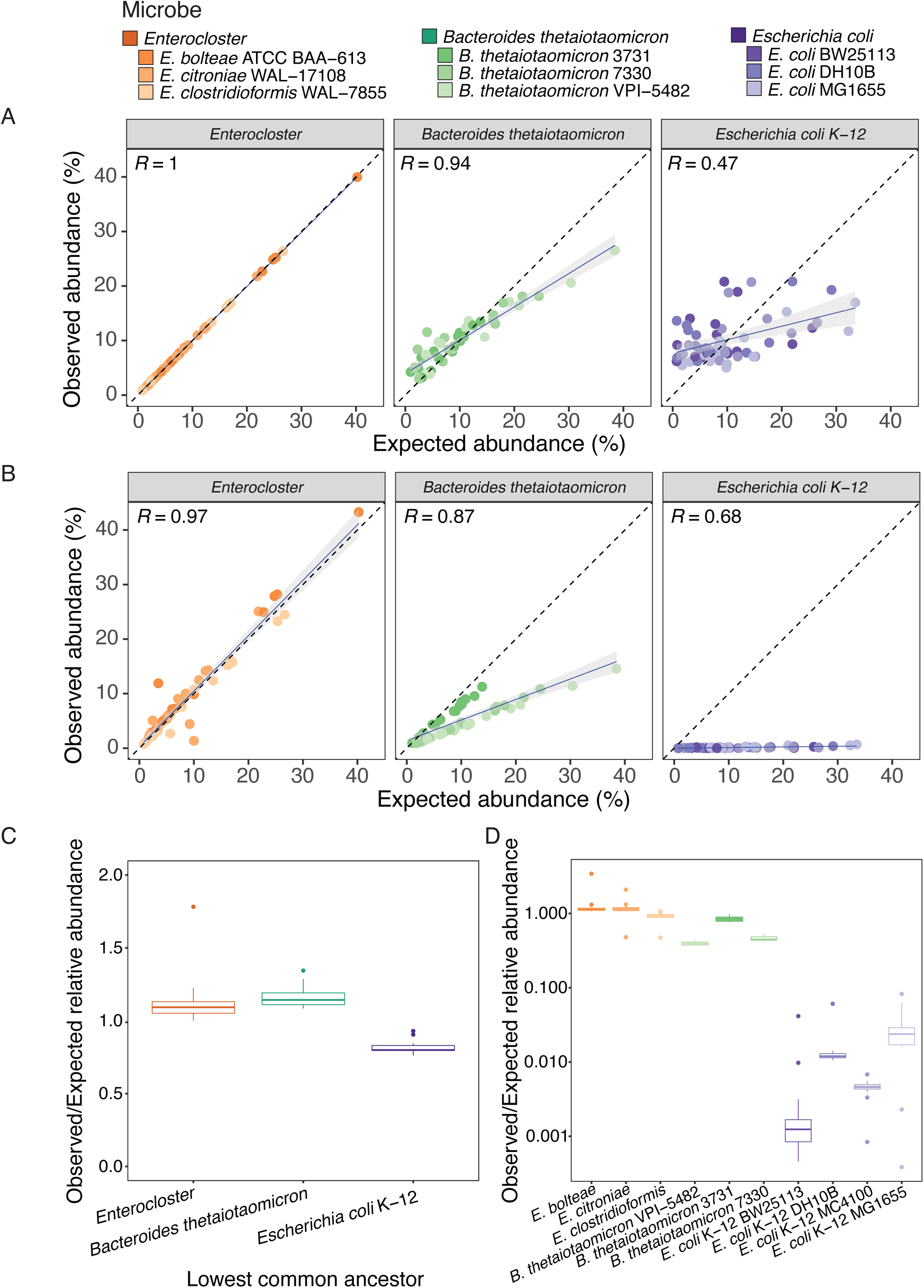
Shotgun sequencing quantification of genetically related microbes by heuristic alignment inaccurately estimates relative abundance due to the random assignment of multimapping reads. (A, B) Scatter plot of observed vs expected relative abundance from *in silico* Illumina reads of the Species-Strain-Substrain (SSS) community members, generated with InSilicoSeq (61). Relative abundance was calculated using a heuristic alignment-based approach with Bowtie2 (44), followed by CoverM (45)(A) and a lowest common ancestor (LCA) approach with Kraken2 (46) (B) to evaluate different shotgun sequencing quantification pipelines (see Methods for details). The dotted line represents a 1:1 slope. The blue line and grey shaded area indicate a linear regression fitted on all data points from each taxon. (C, D) Boxplot of the ratio of observed and expected relative abundance for each LCA (C) and individual member (D) in the SSS community, calculated using the Kraken2 (46) pipeline.

## Discussion

In this study, we introduced PUPpy, a new computational pipeline to design taxon-specific primers in microbial communities (Fig. 1). We benchmarked applications of PUPpy in microbiome research to detect and quantify absolute bacterial counts at high resolution in both defined and complex communities (Fig. 2-4). Our results show that PUPpy-designed taxon-specific primers can detect microbes down to the substrain level in defined bacterial communities (Fig. 2, S1, 3, and S2). Importantly, we also used PUPpy-designed primers to selectively target taxa in complex communities, even when the full membership was unknown (Fig. 3 and S3). Finally, our data suggest that microbe-specific primers can be used in ddPCR to quantify absolute microbial counts, achieving greater accuracy and resolution than short-read 16S rRNA and shotgun sequencing in defined bacterial communities (Fig. 4 and 5).

As a software package, PUPpy was specifically developed to prioritize user-friendliness, with minimal manual handling and command-line knowledge needed. The pipeline can be executed in 2 simple commands either from terminal or a dedicated GUI, taking CDS files as input, and providing a table with taxon-specific primers and key parameters as output. Unlike other primer design tools, PUPpy does not require configuration files, streamlining execution to reduce chances of human error, while also remaining customizable. Users can leverage the flexibility and scalability of PUPpy to design taxon-specific primer sets rapidly and easily for a wide range of custom microbial communities. This is achieved thanks to its alignment-based strategy with MMseqs2 (48), which enables both rapid homology search across all genes in the community and low computational resources, making the pipeline suitable for both computing clusters and personal computers. In the current study, we successfully designed taxon-specific primers on both a computing cluster and personal computer in 2-10 minutes for each 10-member community and under 1 hour for a 167-member complex community on a cluster (see Methods for details). This flexibility allows users to increase, if needed, the number of both targets and non-targets provided to adjust the confidence in primer specificity. In addition, the scalability of PUPpy enables the design of numerous primer sets, also on different genes, for each target, providing multiple ways to confirm specificity in custom microbial communities.

Importantly, increasing the community size, as well as the genetic similarity of its members, makes primer design more challenging. In these communities, the alignment step may not identify any genes sufficiently different to be binned as unique, for instance in the case of *E. coli* MC4100 in the SSS community (Fig. 3B). Users can adjust the alignment stringency, and thus the number of alignments reported, in ‘puppy-align ‘by modifying 4 key parameters: 1) minimum alignment identity, 2) minimum alignment length, 3) minimum alignment coverage, and 4) coverage mode. Lowering the alignment identity, length, and coverage increases stringency by reporting more alignments, which decreases the total number of unique genes found and increases the specificity of taxon-specific primers. Oppositely, raising these values lowers the alignment stringency and increases the number of genes considered to be unique, which may be beneficial in communities where no microbe-specific primers could be designed with default parameters. However, raising the minimum alignment identity, length, and coverage also increases the risk of identifying false positive unique genes. These parameters should thus be manipulated with caution and only if necessary, such as the SSS community which contains, or is expected to contain, multiple genetically related microbes. Despite having relaxed the alignment stringency, PUPpy could not identify truly unique genes for *E. coli* MC4100, but only false positives. This highlights the possibility of achieving greater resolution in shotgun sequencing with appropriate sequencing depth and databases, which may not always be possible with PUPpy-designed PCR primers. Altogether, independently of the manipulation of alignment parameters, experimental validation is essential and always recommended to ensure proper function and specificity of the primers designed.

Beyond primer design in defined communities, we successfully detected a bacterial family with high specificity within a complex microbiota, even without *a priori* knowledge of the microbial membership. This can be a challenging task due to the unknown gene pool, limiting the ability to confirm primer selectivity in the alignment stage. Nevertheless, in this study we were able to design a Muribaculaceae-specific primer cocktail for a conventional mouse gut microbiome by using a comprehensive list of 156 non-targets and 11 intended Muribaculaceae targets. Providing an exhaustive input of non-targets is beneficial to design taxon-specific primers in undefined communities as PUPpy is more likely to identify conserved genes among the target taxa, decreasing the chances of false positive hits even in a complex community. The flexibility and scalability of PUPpy are well suited for this approach, supporting numerous CDS files as input without reaching prohibitive computational requirements. In addition, PUPpy enables users to scale primer design for multiple genes of the same microbial target. This could benefit longitudinal investigations in complex communities where gene transfers, loss or gain could occur. Currently, PUPpy does not directly perform a phylogenetically informed gene choice prior to primer design and instead relies on the non-target list to identify conserved markers among the targets. If needed, users are encouraged to explore phylogenetic tools such as Phylomark (49) or learn about the gene selected by PUPpy prior to their investigations to ensure the selected gene is ideal for the application of interest.

Finally, we have shown the potential of PUPpy-designed microbe-specific primers to accurately quantify absolute microbial counts in defined bacterial communities, even in the presence of genetically related members. Absolute microbial counts can be converted to compositional data, providing comprehensive insights into the dynamics of both individual members and the community. The opposite conversion, from relative to absolute, cannot be performed, confirming the key role of ddPCR and qPCR in quantitative microbial investigations. In these studies, designing and testing primers on multiple genes of the same microbe may be beneficial to ensure genes with a single copy number are being investigated, as these could skew abundance estimation through quantitative PCR approaches. By design, PUPpy does not design primers in genes present in multiple copies when complete microbial assemblies are provided. However, this may fail when scaffold or contig-level assemblies are used as input, as identical genes present more than once may be collapsed to the same location in the genome. Thus, users are encouraged to design multiple primers or confirm copy number prior to investigations focused on highly accurate microbial quantification.

In summary, PUPpy designs taxon-specific primers in diverse microbial communities and represents a fast, user-friendly, and scalable solution to profiling the microbiota. As microbiome research continues to uncover functional diversity beyond the species-level, the ability to detect microbes quickly and accurately below the species level will support high-resolution characterizations of microbial ecosystems that are required for the field to advance.

## MATERIALS AND METHODS

### Resource Table

Resource Table:
| REAGENT or RESOURCE | SOURCE | IDENTIFIER |
| --- | --- | --- |
| **Experimental Models: Organisms** |  |  |
| Swiss-Webster Mice - Germ Free | Taconic, in-house colony | SW GF |
| **Chemicals** |  |  |
| Chopped Meat Medium | Anaerobe Systems | AS-811 |
| BD BBL™ Dehydrated Culture Media: Brain Heart Infusion | Fisher Scientific | B211059 |
| Reinforced Clostridial Medium | Oxoid™ | CM0149B |
| Mucin from porcine stomach (Type III) | Sigma-Aldrich | M1778-100G |
| L-cysteine | Thermo Scientific™ | A1043518 |
| Hemin chloride | MilliporeSigma™ | 37415GM |
| Vitamin K<sub>1</sub> | Alfa Aesar | L10575 |
| Agar | Fisher BioReagents™ | BP1423500 |
| **Critical Commercial Assays** |  |  |
| DNeasy Blood & Tissue Kit | QIAGEN | 69504 |
| DNeasy 96 PowerSoil Pro QIAcube HT Kit | QIAGEN | 47021 |
| Quant-iT™ 1X dsDNA HS Assay | Invitrogen | Q33232 |
| Nextera XT DNA Library Preparation Kit | Illumina | FC-131-1024 |
| 16S rRNA Library Preparation | Sequencing and Bioinformatics Consortium (SBC) | NA |
| 2X DreamTaq Green PCR Master Mix | Fisher Bioreagents | K1082 |
| Agarose | Fisher Bioreagents | BP160-500 |
| ddPCR™ 96-Well Plates | Bio-Rad | 12001925 |
| QX200™ ddPCR™ EvaGreen Supermix | Bio-Rad | 1864034 |
| DG32 Automated Droplet Generator Cartridges | Bio-Rad | 1864108 |
| Pipet Tips for AutoDG™ System | Bio-Rad | 1864120 |
| ddPCR™ Droplet Reader Oil | Bio-Rad | 1863004 |
| Automated Droplet Generation Oil for EvaGreen | Bio-Rad | 1864112 |
| PCR Plate Heat Seal, foil, pierceable | Bio-Rad | 1814040 |
| SYBR™ Safe DNA Gel Stain | Invitrogen | S33102 |
| **Software and Algorithms** |  |  |
| MMseqs2 | https://github.com/soedinglab/MMseqs2 | V14.7e284 (48) |
| R software | https://www.r-project.org/ | Version 4.1.2 (50) |
| ggplot2 | https://www.tidyverse.org/ | Version 3.3.5 (51) |
| tidyverse | https://www.tidyverse.org/ | Version 1.3.1 (52) |
| pyani | https://github.com/widdowquinn/pyani | Version 0.2.9 (53) |
| FastQC | https://www.bioinformatics.babraham.ac.uk/projects/fastqc/ | Version 0.11.9 (54) |
| MultiQC | https://github.com/ewels/MultiQC | Version 1.10.1 (55) |
| fastp | https://github.com/OpenGene/fastp | Version 0.23.2 (56) |
| Bowtie2 | https://github.com/BenLangmead/bowtie2 | Version 2.4.2 (44) |
| CoverM | https://github.com/wwood/CoverM | Version 0.6.1 (45) |
| Kraken2 | https://github.com/DerrickWood/kraken2 | Version 2.1.2 (46) |
| InSilicoSeq | https://github.com/HadrienG/InSilicoSeq | Version 1.5.4 (61) |
| primer3-py | https://github.com/libnano/p | Version 0.6.1 (62) |
|  | rimer3-py |  |
| pandas | https://github.com/pandas-dev/pandas | Version 1.5 (63) |
| biopython | https://github.com/biopython/biopython | Version 1.80 (64) |
| dask | https://github.com/dask/dask | Version 0.15.2 (65) |
| seaborn | https://github.com/mwaskom/seaborn | Version 0.13.0 (66) |
| matplotlib | https://github.com/matplotlib/matplotlib | Version 3.8.2 (67) |
| colorama | https://github.com/tartley/colorama | Version 0.4.1 (68) |
| QIIME2 | https://qiime2.org/ | Version 2023.2 (69) |
| DADA2 | https://github.com/benjjneb/dada2 | Version 1.20 (70) |
| Guppy | https://nanoporetech.com/ | Version 4.5.3 (71) |
| Bonito | https://github.com/nanoporetech/bonito | Version 3.1 (72) |
| Flye | https://github.com/fenderglass/Flye | Version 2.7 (73) |
| Racon | https://github.com/isovic/racon | Version 1.4.22 (74) |
| Medaka | https://github.com/nanoporetech/medaka | Version 1.4.3 (75) |
| CheckM | https://github.com/ECogenomics/CheckM | Version 1.0.18 (76) |
| Prokka | https://github.com/tseemann/prokka | Version 1.14.5 (77) |
| SILVA database | https://www.arb-silva.de/documentation/release-1381/ | Version 138.1 (78) |
| QX Manager 2 | https://www.bio-rad.com/en-ca/life-science/digital-pcr/qx- | Version 2.0 (Bio-Rad) |
|  | software |  |
| **Other** |  |  |
| Standard diet 5K67 - JL Rat & Mouse/Auto 6F autoclavable diet | LabDiet<br>https://www.labdiet.com/product/detail/5k67-labdiet-jl-rat-and-mouse-auto-6f | 5K67 |
| Automated Droplet Generator | Bio-Rad | 1864101 |
| PX1™ PCR Plate Sealer | Bio-Rad | 1814000 |
| C1000 Touch™ Thermal Cycler | Bio-Rad | 1851196 |
| QX200™ Droplet Reader | Bio-Rad | 1864003 |
| **Deposited data** |  |  |
| https://doi.org/10.5683/SP3/HWXPV3 |  |  |

### Ethics statement

All animal experiments were performed according to protocol number (A19-0078) approved by the Animal Care Committee at the University of British Columbia and in direct accordance with the guidelines of the Canadian Council of Animal Care (CCAC).

## EXPERIMENTAL MODELS

### Bacterial strains and culture conditions

Metadata regarding all bacteria used in this study, including taxonomy, sources, and individual culture conditions can be found in Table S1. All bacteria used in this study were cultured anaerobically with the following atmosphere: 5% H_2_, 5% CO_2_, and 90% N_2_ (Linde Canada, Delta, BC, Canada) at 37°C. All media were pre-reduced in an anaerobic chamber (Coy Laboratories, Grass Lake, MI, USA) for at least 24 hours prior to use. Media names, abbreviations, ingredients, and recipes can be found in Table S3. All microbes were streaked onto their respective media plates from glycerol stocks stored at −80°C, followed by inoculation of single colonies into 3 mL of liquid media. Culture conditions and incubation times for individual microbes can be found in Table S1.

### PUPpy Pipeline Implementation

#### Technical specifications

All analyses involving PUPpy were performed on UBC ARC Sockeye, a high-performance computing platform, running an Intel Xeon Silver 4110 (2.1GHz) processor with 16 cores, 192GB DDR4-2666 ECC RAM and 2TB SATA hard disk drive. PUPpy workflows were replicated on a consumer laptop with MacOS Ventura operating system, 10-core CPU with 8 performance cores, 16-core GPU, 16GB RAM and 512 GB solid-state drive to validate the pipeline’s viability under lower computational resources.

### Executing the PUPpy pipeline

The PUPpy pipeline can be executed from terminal by first running ‘puppy-align’, followed by the ‘puppy-primers ‘script. Alternatively, it is possible to execute both commands through a GUI by running ‘puppy-GUÌ and interacting with a button-based system. See below and the PUPpy GitHub repository (https://github.com/Tropini-lab/PUPpy) for more detailed instructions and execution guidelines.

### Sequence alignment of input coding sequence files

The first command of PUPpy, ‘puppy-align’, performs many-against-many local pairwise sequence alignment with MMseqs2 (48) of all user-provided CDSs in a microbial community. This script requires a directory containing the CDS files of the targets for which primers should be designed. Optionally, users can provide a directory with the CDS files of non-targets, which will be exclusively used for specificity checks of the intended targets. Importantly, all CDS files provided as input in the ‘puppy-align’ script will be aligned to ensure specificity of all taxon-specific primers. The actual design of primers occurs in the ‘puppy-primers’ scripts, thus the CDS file choice in ‘puppy-align’ is exclusively meant to indicate which microbes are present in the defined community of interest. CDS files provided by users must meet the following criteria: 1) FASTA headers must be provided in Prokka, RAST or NCBI formats, 2) filenames must contain the string “cds” prior to the extension, and 3) filenames must end with the file extension “.fna”. The command ‘puppy-align’ prepends microbe identifiers prior to the string “_cds” in the CDS filenames to all FASTA headers, which is required for downstream parsing. For improved readability, users are encouraged to name CDS files with easily interpretable and unique microbe identifiers prior to the string “_cds” in the filename. Examples of acceptable CDS filenames include “B_theta_VPI5482_cds.fna” and “B_thetaiotaomicron_VPI_5482_cds_from_genomic.fna”. Following file renaming, PUPpy concatenates all input CDS files (from the primer target and non-target directories) into one file, which is used to create both the query and target databases. Next, PUPpy implements ‘mmseqs search’ from MMSeqs2 (48) (v14.7e284) with default parameters to align the query database against the target database, yielding the output file “ResultDB.tsv” with alignments of every CDS against every other CDS in the input community. The alignment percentage identity threshold of ‘mmseqs search’ can be modified from its default value of ‘-p 0.3’ in ‘puppy-align’ to increase or decrease the alignment stringency. Providing ‘-p’ > 0.3 decreases the alignment stringency and thus increases the chances of finding unique genes. This is not recommended unless the input community is highly related and no unique genes can be found with default parameters, as it will increase the chances of unspecific primer amplification.

### Taxon-specific primer design

The second command of PUPpy, ‘puppy-primers’, designs taxon-specific primers for user-defined primer targets of the input community (Fig. 1A). ‘puppy-primers’ expects the following inputs: 1) the key output file of puppy-align, “ResultDB.tsv”, and 2) a directory containing the CDS files of the primer targets PUPpy should design taxon-specific primers for (Fig. 1A). As default, ‘puppy-primers’ designs microbe-specific primers (Fig. 1B). For example, given a 3-member bacterial community composed of *Bacteroides thetaiotaomicron*, *Bacteroides ovatus*, and *Escherichia coli*, a microbe-specific primer set would only amplify *B. thetaiotaomicron* but no other microbe. Users can request PUPpy to design group-specific primers by adding the flag ‘-p group’ (Fig. 1B). For example, given the same 3-member community above, a *Bacteroides*-specific group primer set would amplify both *B. ovatus* and *B. thetaiotaomicron*, but not *E. coli*. Depending on the mode, ‘puppy-primers’ parses the alignment file to identify candidate CDSs to design microbe- or group-specific primers. CDSs that only align to themselves are considered for microbe-specific primers, while CDSs present in all intended targets are considered for group-specific primers. ‘puppy-primers’ orders candidate genes in decreasing length and then utilizes Primer3 (62) to design primers, while giving flexibility to adjust key primer design parameters either through terminal or GUI. A comprehensive list of all modifiable Primer3 primer design parameters, including all default values, is available by running ‘puppy-primers -h’. The key output of PUPpy ‘puppy-primers’ for microbe- and group-specific primers are “UniquePrimerTable.tsv” and “GroupPrimerTable.tsv”, respectively. Both files contain information about the identified gene name, gene sequence, primer pair penalty scores, amplicon size, forward and reverse primer sequence, length, GC content and melting temperature for all the user-defined primer targets. Running ‘puppy-primers’ on microbe-specific mode also yields the barplot “UniqueGenesPlot.pdf”, which shows the number of unique and non-unique genes found for every primer target of the input community.

### DNA Extraction and Quantification

Bacterial gDNA was extracted using the DNeasy Blood & Tissue Kit (QIAGEN) and quantified with 3 technical replicates using the Quant-iT™ 1X dsDNA HS (High-Sensitivity) Assay (Invitrogen), following the manufacturer’s instructions. Microbial gDNA measured concentrations in ng/μL were converted to copies/μL using the formula below, where *X* is the amount of DNA in ng/μL, and *N* is the genome size in base pairs (bp) as reported by the NCBI (Table S1).

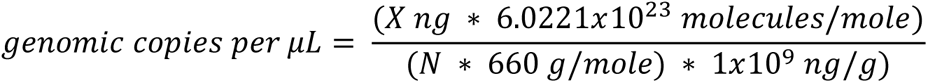

## Primer design and specificity validation

All taxon-specific primers used in this study were designed with PUPpy (see Table S2) and ordered with Thermo Fisher Scientific (https://www.thermofisher.com/order/custom-standard-oligo/). Custom Standard DNA Oligos were ordered dry, desalted, and with a synthesis scale of at least 25 nmole.

### GUT and SSS defined communities

Taxon-specific primers, and their respective key parameters, designed for the GUT and SSS communities can be found in Table S2. Microbe-specific primers for both the GUT and SSS community were designed separately by running ‘puppy-align’ with default values, followed by ‘puppy-primers’ with the parameters ‘-s 125 175 -optm 62 -tmd 1.5’. Group-specific primers targeting the 3 major taxa of the SSS community were designed in 3 distinct runs by running ‘puppy-align’ with default values, followed by ‘puppy-primers’ with the parameters ‘-p group -pr <TAXON_CDSS> -s 125 175 -optm 62 -tmd 1.5’, where *taxon_CDSs* were the CDS files of the1) *Enterocloster* species, 2) *B. thetaiotaomicron* strains, and 3) *E. coli* substrains. Primer specificity was experimentally validated by PCR and gel electrophoresis. Each taxon-specific primer in the GUT and SSS communities was validated against 4 conditions: 1) the gDNA(s) targeted by the primer pair 2) the 10-member pool gDNA, 3) the 10-member pool without the target gDNA(s), and 4) a no-template control (water). The 10-member GUT and SSS communities were made by pooling an equal ratio of genomic copies of the respective microbial gDNA samples (quantified as above).

### Muribaculaceae complex community

The Muribaculaceae-specific primer cocktail was created by pooling microbe-specific primer pairs targeting individual *Muribaculum* members (see Table S2). To maximise the chances of specificity within a complex community, microbe-specific primers targeting *Muribaculum* members were designed against a comprehensive input community of 167 gut commensals (including the target Muribaculaceae) available in the Tropini Strain Library (see Table S1). Although all Muribaculaceae-specific primers target the same CDS across intended targets, distinct primers were designed to account for base pair differences that may have affected primer annealing. PUPpy was used to identify a gene shared across all Muribaculaceae members and individual primers were manually designed to maximise specificity across all target members. The specificity of the Muribaculaceae-specific primer cocktail within a complex community was validated using a Muribaculaceae-depleted and -positive conventional mouse model (42). Conventional mice were treated with 15% (w/v) osmotic laxative polyethylene glycol (PEG) for 7 days. Following PEG administration, fecal samples from both treated (Muribaculaceae-depleted) and untreated (Muribaculaceae-positive) mice were collected and total DNA was extracted from faecal contents using DNeasy 96 PowerSoil Pro QIAcube HT Kit (QIAGEN, Germantown, MD, USA), following the manufacturer’s protocol. Muribaculaceae presence or absence in both samples was confirmed by 16S rRNA sequencing. The Muribaculaceae-specific primer cocktail was experimentally validated by PCR and gel electrophoresis against the following 4 conditions: i) 11 distinct *Muribaculum* gDNA samples, ii) no-template control (water), iii) Muribaculaceae-depleted samples, and iv) Muribaculaceae-positive.

All PCR reactions used to experimentally validate primers contained 7.5 μL of 2X DreamTaq Green PCR Master Mix (Thermo Fisher^TM^), 5 μL of NF-H_2_O, 0.75 μL of forward and reverse primers (final concentration of 340 nM) and 1 μL of DNA template. PCR amplification was performed on a C1000 Touch^TM^ Thermal Cycler (Bio-Rad) with the following program: 98°C for 2 min, 40 cycles of (98°C for 30 s, *specific_annealing_T* for 30 s, 72°C for 15 s), 72°C for 5 min and 4°C hold. The annealing temperature of each primer pair (*specific_annealing_T)* can be found in Table S2. Extension time was decreased from 1 minute to 30 seconds to avoid unspecific amplification. Following amplification, 8 μL of PCR product was run on a 1.5% agarose gel stained with SYBR^TM^ Safe DNA Gel Stain (Invitrogen) at 90V for 60 minutes. Primer specificity was assessed by visually inspecting gels for the presence or absence of bands (Fig. S1, S2 and S3).

### Shotgun and 16S rRNA Sequencing and Analyses

Library preparation and sequencing for the defined communities were performed at Sequencing + Bioinformatics Consortium (SBC) at the University of British Columbia (Vancouver, Canada). Extracted DNA was quantified using Qubit fluorometry. Shotgun metagenomic libraries were then prepared using the Illumina DNA prep library preparation kit (Illumina, San Diego, CA, USA). These libraries were sequenced on a NextSeq Mid output flow cell, generating paired-end 150 bp reads. For 16S, the V3 and V4 regions of the 16S rRNA gene were PCR amplified using the following primers (79): F: 5′-TCGTCGGCAGCGTCAGATGTGTATAAGAGACAGCCTACGGGNGGCWGCAG and R: 5′-GTCTCGTGGGCTCGGAGATGTGTATAAGAGACAGGACTACHVGGGTATCTAATCC. These amplicons were then converted to sequencing libraries using an 8-cycle indexing PCR with Nextera XT primers (Illumina, San Diego, CA, USA). These libraries were sequenced on a MiSeq v2 flow cell (Illumina, San Diego, CA, USA) to generate paired-end 250 bp reads. Raw base call data (bcl) were converted into fastq format using the bcl2fastq conversion software from Illumina. All computational analyses were performed on the UBC ARC Sockeye high-performance computing platform. 16S library preparation (80) for the Muribaculaceae-depleted and -positive samples was performed at Gut4Health, BC Children’s Hospital Research Institute, Vancouver, BC, Canada. The V4 region was amplified using the following sequences: F: 5′-TATGGTAATTGTGTGYCAGCMGCCGCGGTAA and R: 5′-AGTCAGTCAGCCGGACTACNVGGGTWTCTAAT. Amplicon libraries were purified, normalized, and pooled using the SequalPrep^TM^ normalization plate (Applied Biosystems). Library concentrations were verified using a Qubit TM dsDNA high sensitivity assay kit (Invitrogen) and KAPA Library Quantification Kit (Roche) following manufacturer details. The purified pooled libraries were submitted to the Bioinformatics + Sequencing Consortium at UBC which verifies the DNA quality and quantity using an Agilent high sensitivity DNA kit (Agilent) on an Agilent 2100 Bioanalyzer. Sequencing was performed on the Illumina MiSeq^TM^ v2 platform with 2 x 250 paired end-read chemistry.

For shotgun sequencing, the quality of raw reads was evaluated using FastQC (54) (v0.11.9) and MultiQC (55) (v1.10.1). Raw reads were cleaned using fastp (56) (v0.23.2) with the parameters ‘-q 20 -p - 3 −5 -M 20 -W 4 -w 15 -c -l 50 --dedup --dup_calc_accuracy 5’. Cleaned reads were used to calculate relative abundance using both a heuristic alignment approach and an LCA approach. In the heuristic alignment-based approach, reads were mapped to a custom reference index of the 10 concatenated microbial genomes using Bowtie2 (44) (v2.4.2) with the following parameters: ‘*-D 20 -R 3 -N 0 -L 20 -i S,1,0.50 --reorder --no-unal -p 40’.* Relative abundance of individual microbes was calculated using CoverM (45) (v0.6.1) with the following key parameters ‘coverm genome -m relative abundancè. In the LCA approach, relative abundance was calculated with Kraken2 (46) (v2.1.2) using trimmed reads (as described above) and a custom database of the 10 respective community members.

For 16S rRNA sequencing, raw reads were quality checked with FastQC (54) (v0.11.9) and then analysed with QIIME2 (69) (v2023.2) to estimate microbial relative abundance. Reads were processed with DADA2 (70) (v1.20) to trim primers and truncate sequences with quality score below 30. Denoised sequences were assigned taxonomic labels using a classifier trained on the SILVA (78) (v138.1) 99% database.

### *In silico* Illumina reads generation and data analysis

*In silico* Illumina reads for the SSS community were simulated with InSilicoSeq (61) (v1.5.4) to achieve known input proportions, using the following parameters: ‘--mode basic --draft <SSSGENOMES> -- coverage <INPUTCOVERAGE> --n_reads 1.1M’. The genomes of all SSS members were used to generate *in silico* reads at desired input proportions, determined by coverage. To achieve a wide range of input coverages for each member, all 5 default ‘--coveragè values offered by InSilicoSeq (61) were used: uniform, halfnormal, lognormal, exponential, and zero_inflated_lognormal. Relative abundance from these *in silico* Illumina shotgun sequencing reads was calculated using both heuristic alignment and LCA approaches, as described above.

### Quantification of Absolute Microbial Counts

Absolute quantification of microbial counts, measured in copies/μL, for both the 10-member GUT and SSS pools was achieved with droplet digital PCR (ddPCR). All ddPCR reactions were run at Gut4Health (RRID:SCR_023673). The same microbe-specific primers validated for the GUT and SSS communities were used in ddPCR (see Table S2). The respective 9-member pools for each primer pair, lacking the intended target, were run as negative controls. Absolute microbial counts were converted to relative abundance by dividing the copies/μL of each organism by the sum of copies/μL of all organisms in the community. Microbial relative abundance measured by ddPCR in the SSS community was calculated from 9 organisms instead of 10 as we could not design microbe-specific primers for *E. coli* MC4100.

Briefly, the 10-member GUT and SSS gDNA pools were diluted in NF-H_2_O (Fisher Bioreagents) to achieve an expected concentration of the target DNA between 0 and 5000 copies/μL. Every ddPCR reaction was prepared in a semi-skirted ddPCR 96-well plate (Bio-Rad) and contained 11 μL 2X QX200^TM^ ddPCR^TM^ EvaGreen Supermix (Bio-Rad), 0.45 μL of forward and reverse primers (final concentration of 204 nM), 8.1 μL of NF-H_2_O, and 2 μL of the diluted gDNA pool. The ddPCR plate was placed in an Automated Droplet Generator (Bio-Rad), together with DG32 Automated Droplet Generator Cartridges (Bio-Rad), Pipet Tips for AutoDG^TM^ system (Bio-Rad), and Automated Droplet Generation Oil for EvaGreen (Bio-Rad). Following droplet generation, the ddPCR plate was sealed with pierceable foil (Bio-Rad) using a PX1^TM^ PCR Plate Sealer (Bio-Rad). After sealing, the ddPCR plate was placed in a C1000 Touch^TM^ Thermal Cycler (Bio-Rad) and amplified using the following program: 95°C for 5 min, 40 cycles of (95°C for 30 s, 62°C for 1 min), 4°C for 5 min, 90°C for 5 min and 4°C hold. All changes in temperature were limited to 2°C/s ramp rate. Droplet reading was performed with a QX200^TM^ Droplet Reader (Bio-Rad) using ddPCR^TM^ Droplet Reader Oil (Bio-Rad). Manual selection of the amplitude threshold and data analysis were performed on the QX Manager v2.0 software (Bio-Rad).

### Average Nucleotide Identity Analyses

Average nucleotide identity (ANI) analyses for the GUT and SSS communities were performed using the ANIb method of pyani (53) (v0.2.9). The CDS files of *E. coli* BW25113, MG1655, and DH10B were concatenated with the ‘cat’ command and provided as pyani input (-i flag) together with the *E. coli* MC4100 CDS file. The ANI outputs were imported into RStudio and heatmaps were plotted using ggplot2.

### *M. intestinale* G6 sequencing and assembly

#### High molecular weight genomic DNA extraction

*M. intestinale* G6 was cultured following the conditions outlined in Table S1. High molecular weight genomic DNA (gDNA) was isolated using a Genomic-tip 100/G Kit (QIAGEN) following the manufacturer’s protocol. Extracted DNA was allowed to relax at 4°C overnight and was then quantified using a Quant-iT™ 1X dsDNA HS Assay Kit (Invitrogen) and quality controlled using spectrophotometry. DNA integrity was confirmed on a 1% agarose gel.

### Genome sequencing

Library construction on gDNA extracted from pure *M. intestinale* G6 cultures was performed with NEBNext® Companion Module for Oxford Nanopore Technologies® (ONT) Ligation Sequencing (New England Biolabs, MA, USA), ONT Ligation Sequencing Kit (SQK-LSK109, Oxford Nanopore Technologies, UK) and Flongle Sequencing Expansion kit (EXP-FSE001, Oxford Nanopore Technologies, UK) following the manufacturer’s instructions. Sequencing was performed with Flongle flowcell (FLO-FLG001) on MinION sequencer (ONT).

### *De novo* genome assembly

Sequencing raw reads were basecalled with Guppy (71) (v4.5.3) and Bonito (72) (v3.1) model and then assembled with Flye (73) (v2.7) to generate a draft genome. The draft genome was first polished with Racon (74) (v1.4.22) using -m 8 -x −6 -g −8 -w 500, then polished with Medaka (75) (v1.4.3) using Medaka model for Bonito.

The final assembly resulted in a single 3,123,480 bp circular genome, with average coverage of 467x. The assembly quality was evaluated using CheckM (76) (v1.0.18), which showed 98.4% completeness and 0.26% contamination. The *de novo* genome assembly was annotated using Prokka(77) (v.1.14.5) using the following settings: I) --addgenes, II) --addmrna III) --gffver 3 IV) --genus Muribaculum, V) -- kingdom Bacteria, VI) --gcode 11 which identified including 2918 CDSs.

## Data and Software Availability

Code, including vignettes and tutorials, are available in the PUPpy GitHub page: https://github.com/Tropini-lab/PUPpy

Data is available at: https://borealisdata.ca/privateurl.xhtml?token=37d3e60b-62ca-40d8-9696-f77f9d7ccb57

## Author contributions

H.G., Y.M.F., J.C.B., K.M.N., and C.T.: conceptualization, methodology, and software. H.G., Y.M.F., J.C.B., K.M.N., D.M.P., and C.T.: resources. H.G., C.T.: writing. H.G., Y.M.F., D.M.P.: formal analysis, validation, investigation, data curation, and visualization. C.T.: supervision, project administration, and funding acquisition. All authors read and approved the paper prior to submission.

## Acknowledgments

The authors acknowledge that the land we performed this research on is the traditional, ancestral, and unceded territory of the xwməθkwəy̓əm (Musqueam) Nation. We encourage others to learn more about the native lands in which they live and work at https://native-land.ca/

The authors would like to thank Ian Ghezzi, Paula Otto Andrade von Sperling, Bryan Merrill, Giselle McCallum, Jerry He and Thad Hughes for useful discussions, Claudia Stroppa for designing the PUPpy logo, Jerry He for inspiration on the “PUPpy” name, Sam Collins for support with mouse experiments, and Giselle McCallum for reading the manuscript and providing critical feedback. We also thank Angele Arrieta for research support. This work received support from Gut4Health (RRID:SCR_023673) at the BC Children’s hospital Research Institute, and the Sequencing + Bioinformatics Consortium (SBC) and Advanced Research Computing (ARC) at the University of British Columbia. The authors acknowledge support from 4-Year Fellowship (to H.G., J.C.B, and D.M.P.), NSERC PGS-D Scholarship (to D.M.P.), Canadian Institute for Advanced Research / Humans and the Microbiome (FL-001253 Appt 3362, to C.T.), Michael Smith Foundation for Health Research Scholar Award (18239, to C.T.), Johnson & Johnson Women in STEM2D Award (015007, to C.T.), and Canada Foundation for Innovation / Infrastructure Operating Fund (38277).

---

**<u>Fig. S1</u>**

PUPpy-designed microbe-specific primers selectively amplify all targets in the GUT community. Related to Fig. 2. Experimental validation of all microbe-specific primers by polymerase chain reaction (PCR) and gel electrophoresis, following the conditions outlined in Fig. 2C. All 10 microbe-specific primers selectively amplify their respective target, showing no unintended amplification. Faint primer dimers are visible in *A. muciniphila*, *B. thetaiotaomicron*, *C. sporogenes*, *E. faecalis*, and *E. rectale*.

**<u>Fig. S2</u>**

PUPpy-designed microbe- and group-specific primers selectively amplify the respective targets in the SSS community. Related to Fig. 3. (A) Experimental validation of all microbe-specific primers by polymerase chain reaction (PCR) and gel electrophoresis, following the conditions outlined in Fig. 3C. The initial validation of *E. clostridioformis* specific primers (top right gel) against the (-) pool without the intended target (condition *iii*) showed unspecific amplification at different sizes than the expected amplicon. Decreasing the extension time to 30 seconds (instead of 1 minute) prevented unspecific amplification, as seen in the small gel. The water control was not repeated because condition iii also acts as a negative control. (B) Experimental validation of group-specific primers by PCR, following the conditions outlined in (A). The gels were imaged on distinct runs. All group-specific primers selectively amplified their respective targets without unintended or unspecific amplification.

**<u>Fig. S3</u>**

PUPpy-designed Muribaculaceae-specific primers selectively amplify Muribaculaceae members in a complex microbial community. Related to Fig. 3E and 3F. Experimental validation of Muribaculaceae-specific primers by polymerase chain reaction (PCR) and gel electrophoresis, following the conditions outlined in Fig. 3E. The 2 PCR gel images were taken in distinct instances. The Muribaculaceae-specific primer mix amplified 10 different Muribaculaceae members, including strains that were not originally included in the PUPpy input for specificity checks (See Table S1 and S2). *M. intestinale* NM03 was run twice. *M. intestinale* NM65 was the only member not to be amplified, likely due to low DNA content. In the fecal samples, the Muribaculaceae-specific primer mix accurately discriminated the absence (*iii*) and presence (*iv*) of Muribaculaceae members in a complex microbial community, while also enabling strain-level resolution (*i*). The numbers in brackets for the fecal samples indicate biological replicates.

**<u>Fig. S4</u>**

Microbial quantification using only strain-informative shotgun sequencing reads under-estimates abundance of strains and substrains. Related to Figure 4C. Microbial quantification of SSS community members was estimated using shotgun sequencing (left panel) and ddPCR (right panel). In shotgun sequencing, relative abundance of the total reads was quantified using Bracken, Kraken2 at the Lowest Common Ancestor (LCA) level, and Kraken 2 while only considering unique reads, which are reads that map unambiguously to individual members of the community.

## Table legends

**Table S1. Metadata of all microbes used for the design and validation of PUPpy-designed taxon-specific primers.** Microbes with “NA” noted under the growth conditions columns (last 4) were not cultured but only used for *in silico* primer design. Media ingredients and recipes for culture conditions can be found in Table S3.

**Table S2. Primers sequences and essential parameters for taxon-specific primers designed and validated in this study.** The column ‘target organisms’ exclusively refers to the specificity within the community for which primers were designed and does not necessarily extend to other microbial communities.

**Table S3. Media ingredients and recipes used to culture microbes in this study.** Details on culture conditions and media for each microbe can be found in Table S1.

