## Supplementary figures and images for "PUPpy: a primer design pipeline for substrain-level microbial detection and absolute quantification"

### Figure S1

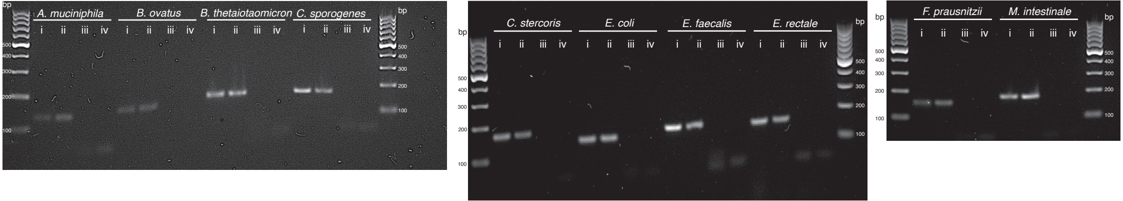

### Figure S2

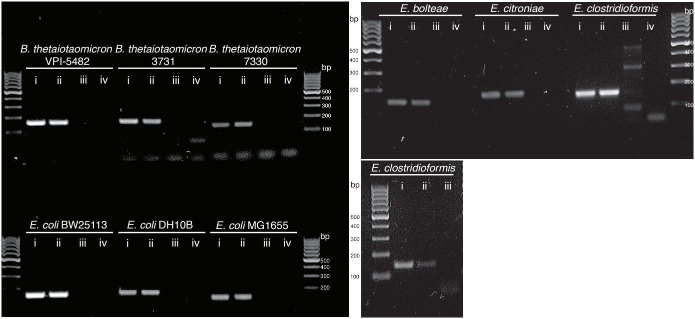

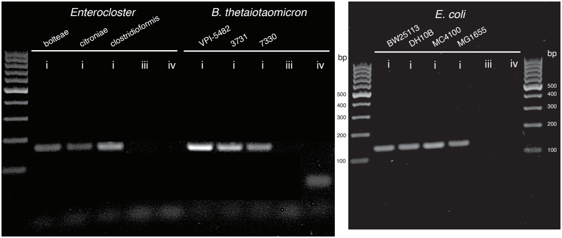

### Figure S3

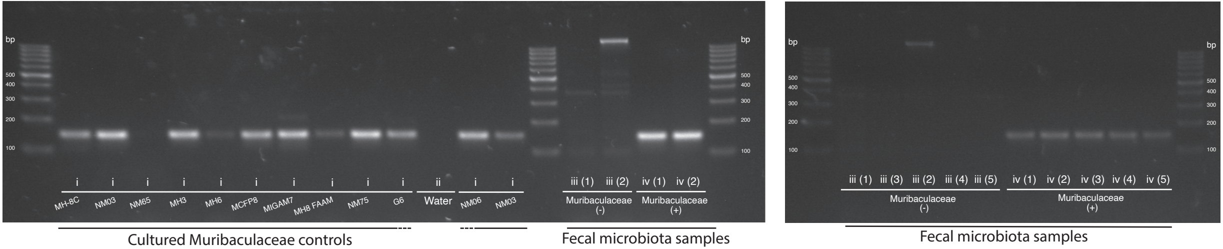

### Figure S4

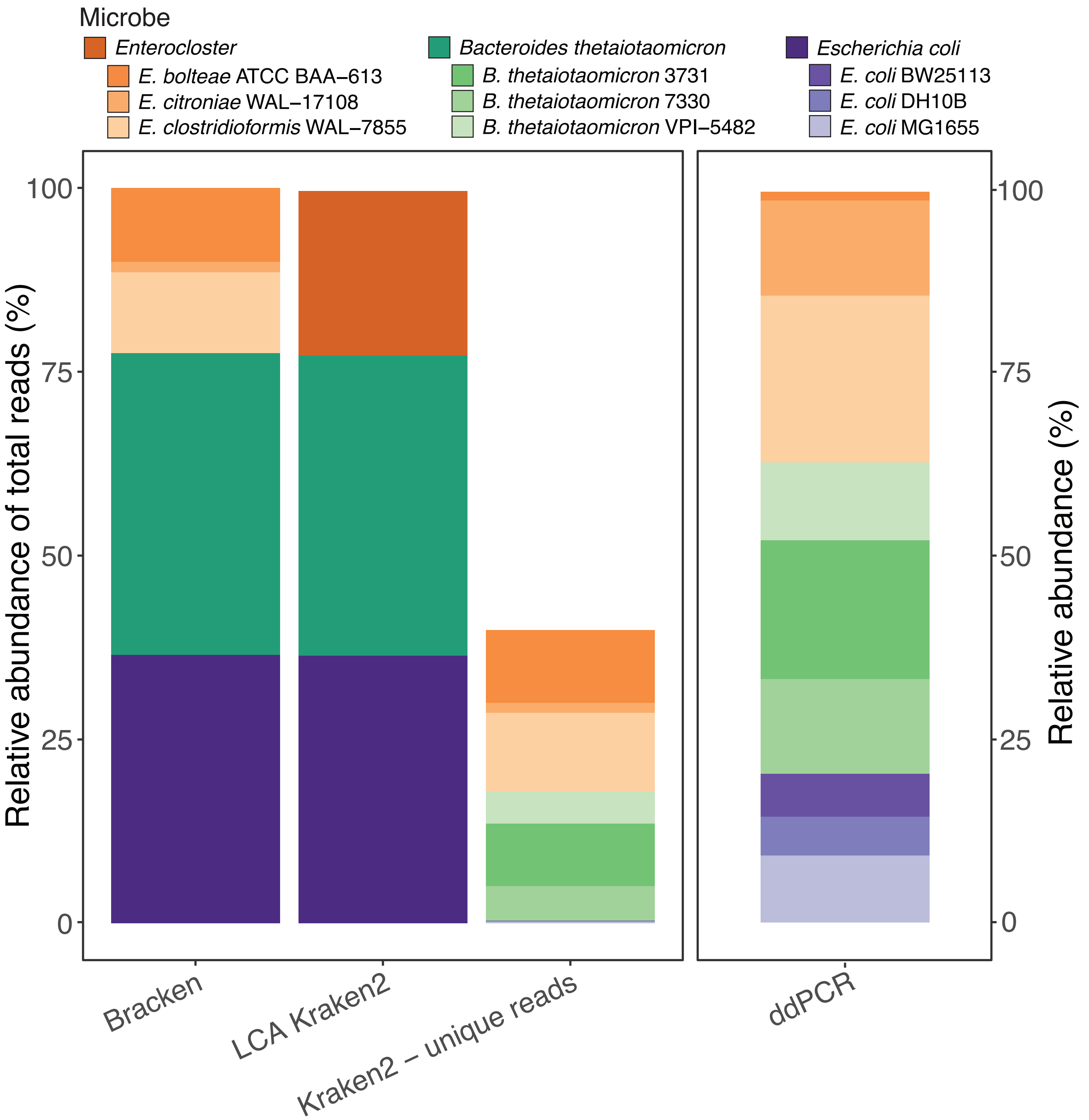
