## Supplementary material for "PUPpy: a primer design pipeline for substrain-level microbial detection and absolute quantification": Table S3

| **Medium** | **Abbreviation** | **Ingredients (per Liter)** | **Recipe** |
| --- | --- | --- | --- |
| Brain heart infusion-supplemented^1^ | BHIS | 37 g Bacto^TM^ Brain Heart Infusion (BHI) (BD Biosciences, San Jose, CA, USA) and (for plates) 15 g agar.  Filter-sterilised supplements:  - 1.5 mL of 333 mg/mL L-cysteine hydrochloride (5 g L-cysteine dissolved in 15 mL 1 N HCl),  - 2 mL of 5mg/mL hemin (50 mg hemin dissolved in 1 mL 1 N NaOH and 10 mL ddH2O)  - 1 mL of 1 mg/mL Vitamin K3 (100 μL Vitamin K3 in 10 mL 95% ethanol) | - Suspend BHI and (optional) agar in MilliQ water  - Autoclave at 121°C for 15 minutes  - When cooled to ~55°C add supplements to the medium and stir gently  - Pour plates if doing solid media |
| Brain heart infusion-supplemented plus 0.5% mucin^1^ | BHIS+0.5% muc | 37 g Bacto^TM^ Brain Heart Infusion (BHI), 5 g porcine mucin (Sigma Aldrich, St. Louis, MO, USA), and (for plates) 15 g agar.  Filter-sterilised supplements as per BHIS medium | - Suspend BHI, mucin, and (optional) agar in MilliQ water  - Autoclave at 121°C for 15 minutes  - When cooled to ~55°C add supplements to the medium and stir gently  - Pour plates if doing solid media |
| Reinforced clostridial medium | RCM | 38 g Oxoid^TM^ Reinforced Clostridial Medium (RCM) (Thermo Fisher Scientific, Waltham, MA, USA) and (optional) 15g agar. | - Suspend RCM and agar in MilliQ water  - Autoclave at 121°C for 15 minutes |
| Chopped Meat Broth | CM | Chopped Meat (CM) Broth (Anaerobe Systems, Morgan Hill, CA, USA) | NA |

**Table S3.** Media ingredients and recipes used in this study.
